# Curative levels of endogenous gene replacement achieved in non-human primate liver using programmable genomic integration

**DOI:** 10.1101/2024.10.12.617700

**Authors:** Jenny Xie, Maike Thamsen Dunyak, Patrick Hanna, Angela X. Nan, Brett Estes, Jesse C. Cochrane, Shuai Wu, Jie Wang, Connor McGinnis, Qiang Wang, Rejina Pokharel, Dev Paudel, Jason Zhang, Dan Li, Parth Amin, Siddharth Narayan, Angela Hsia, Dane Z. Hazelbaker, Xiarong Shi, Meredith Packer, Brian Duke, Ryan Dickerson, Charlotte Piard, Martin Meagher, Jason Gatlin, Sonke Svenson, Adrianne Monsef, Raymond W. Bourdeau, Kieu Lam, Steve Reid, Mohammad Kazemian, Nisher Chander, Richard Holland, James Heyes, Swati Mukherjee, Sandeep Kumar, Daniel J. O’Connell, Jonathan D. Finn

**Affiliations:** Tome Biosciences, Watertown, MA, USA

## Abstract

The ability to efficiently place a large piece of DNA in a specific genomic location has been a goal for the gene therapy field since its inception; however, despite significant advances in gene editing technology, this had yet to be achieved. Here we describe two methods of programmable genomic integration (PGI) that overcome some of the limitations of current approaches. Using a combination of clinically validated delivery technologies (LNP, AAV), we demonstrate the ability to specifically integrate large (>2 kb) DNA sequences into endogenous introns in the liver of non-human primates (NHP). PGI was effective across multiple genomic locations and transgenes, and insertion led to expression from the endogenous promoter. PGI was highly efficient, achieving expression in >50% of liver cells after a single course of treatment, which would be curative for most monogenic recessive liver diseases. This is the first report of clinically curative level of gene insertion at endogenous loci in NHP.

## Introduction

The ability to replace defective genes at their endogenous locations would address many limitations of current gene therapy and gene editing systems. It would enable native regulation, (avoiding over or under-expression), a single product that will treat most if not all patients (independent of mutation), and allow for a one-time cure that adapts as the patient grows.

To date, *in vivo* gene insertion approaches have primarily focused on random/semi-random integration^1, 2^ or nuclease-based capture of AAV genomes^3–5^. These approaches are limited either by low efficiency and require integration into highly expressed genomic ‘safe harbors’ (e.g. Albumin) or require small, non-native promoters. None of these approaches can maintain endogenous expression or native regulation of the integrated gene, which is a critical limitation given the importance of dynamic gene regulation in virtually all aspects of life.

Programmable Genomic Integration (PGI) is the ability to insert a desired DNA sequence into a user defined genomic location. Integrase-mediated Programmable Genomic Integration (I-PGI) allows for the insertion of large DNA sequences into specific genomic locations, enabling the efficient replacement of defective genes at their native loci. I-PGI, innovated from PASTE^6^, uses a Cas9 nickase, a writing enzyme (e.g. reverse transcriptase), and attachment guide RNAs (atgRNAs) to write a large serine integrase (LSI) landing site, herein referred to as a ‘beacon’ (∼40-50 bp, attB or attP) at a specific location in the genome. Expression of the corresponding LSI, along with a template DNA containing the cognate recognition site, leads to the targeted integration of the template DNA at the beacon (Fig. 1a, Extended data Fig. 1a-c). LSIs are agnostic to the size of the template DNA and their recombination is irreversible, unidirectional and independent of host cell DNA repair mechanisms allowing activity in all cell types, both dividing and quiescent.

**Figure 1.**
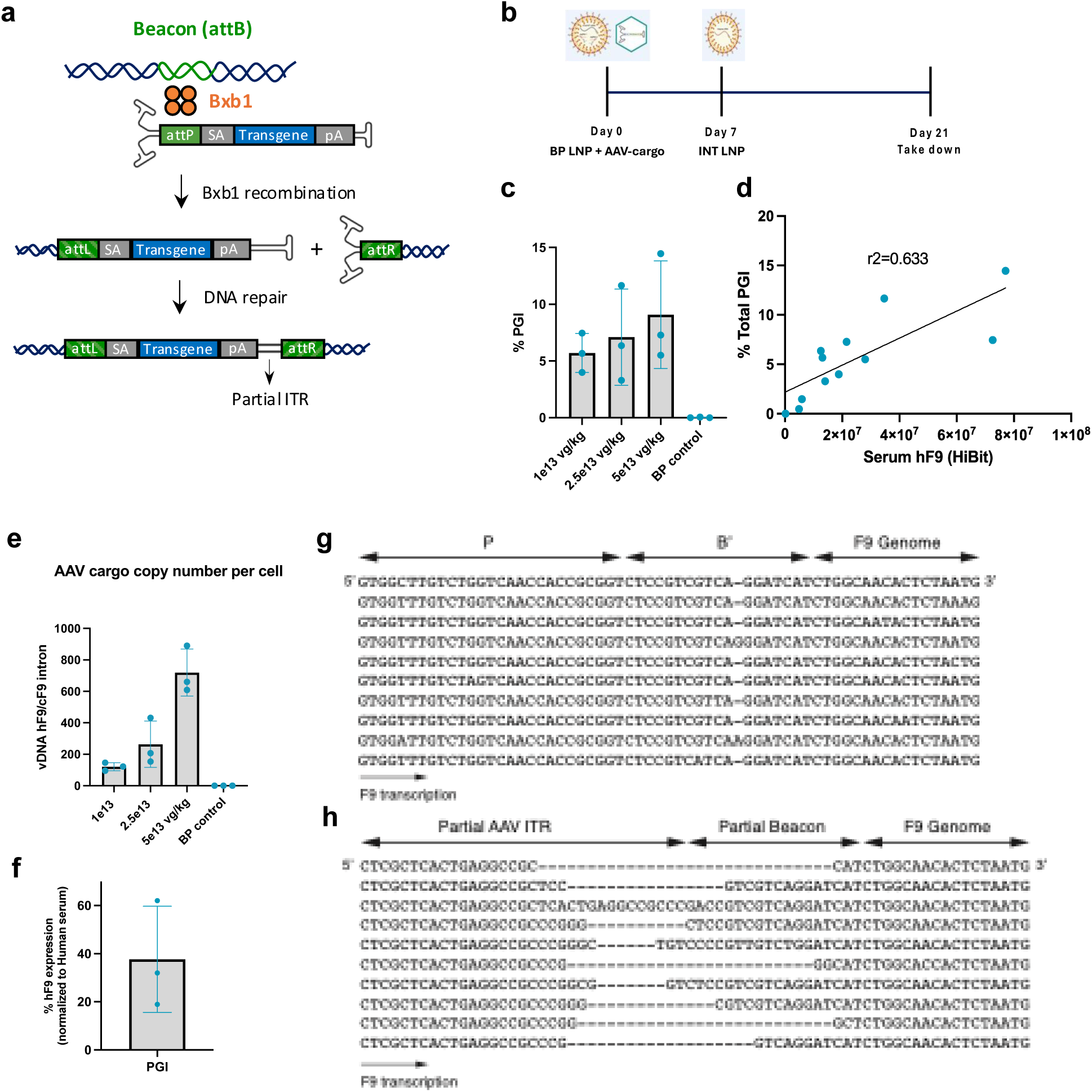
Study design and edit outcomes for programmable genomic integration (PGI) in non-human primates (NHP) (**a**) Diagram of AAV attP cargo integration into placed attB (beacon) in host genome by Bxb1 integrase showing genomic outcome after linear genome incorporation. (**b**) Dosing regimen showing timing of AAV cargo, BP LNP containing nCas9-RT mRNA, atgRNA 1, atgRNA 2, and INT LNP containing integrase mRNA, and takedown. (**c**) Quantitation of total human F9 (hF9) gene insertion into cynomolgus F9 (cF9) in NHP at 3 AAV doses showing increase in insertion frequency with higher cargo doses. (**d**) Correlation between insertion rate determined by ddPCR and relative hF9 levels in serum from insertion of the HiBit tagged gene determined by luminescence reporter assay. (**e**) Relative AAV vector genome copy number in bulk NHP DNA extract. ddPCR was used to quantify AAV vector genome copy and NHP genome copy and results are reported as the ratio of the two measurements for each animal. (**f**) hF9 expression level in NHP after PGI. hF9 was quantified by ELISA in NHP serum using pooled human serum from healthy donors as a standard. (**g**) Multiple sequences alignments (MSA) of I-PGI junction sequences exhibits a high degree of uniformity with minimal variation in the P and B’ sequences. (**h**) MSA of the DF-PGI junction sequences shows high sequence diversity with variations in the inverted terminal repeat (ITR) regions and partial beacon sequences.

Lipid nanoparticles (LNPs) and adeno-associated viruses (AAVs) are clinically relevant, highly effective delivery systems. A single administration of LNP carrying Cas9 mRNA and a chemically synthesized sgRNA achieved >90% knockdown of TTR protein in patients with transthyretin amyloidosis (ATTR)^7^. While LNPs can efficiently deliver cargo to the cytoplasm and minimize the immunogenicity and off-target risks associated with long-term expression of editing enzymes, they are not able to directly access the nucleus. In contrast, AAVs have evolved specifically to deliver DNA cargo into the nucleus and multiple serotypes have been clinically validated for hepatocytes^8^. For our *in vivo* PGI studies, we have employed LNPs for the transient delivery of editing machinery and AAV for nuclear delivery of the template DNA. Here, we describe our progress in developing PGI for the targeted insertion of genes at their endogenous locations in liver at efficiencies considered to be clinically curative.

## Results and Discussion

Using Hemophilia B and Factor 9 (F9) as proof-of-concept, we achieved therapeutically relevant levels of PGI (>10%) in bulk liver of non-human primates (NHPs) using a combination of LNP and AAV delivery. Specifically, we co-dosed a beacon placement LNP (BP-LNP) at 3 mg/kg containing fused Cas9 nickase (nCas9) and engineered reverse transcriptase mRNA, along with a pair of attachment guide RNAs (atgRNAs) targeting intron 1 of the cyno F9 (cF9) locus and AAV at increasing doses of 1e13 - 5e13 vg/kg encoding a promoterless cargo consisting of Bxb1 attP, engineered splice acceptor, codon-optimized human F9 (hF9) exons 2-8, and C-terminal HiBit tag. The integrase LNP (INT-LNP) was dosed 1 week later at 3 mg/kg and animals were sacrificed after an additional 14 days at Day 21 (Fig. 1b). Integration in bulk liver tissue was assessed by ddPCR junction assays targeting the attL and attR recombination sites in cF9 intron 1. We observed a modest dose response with increasing amount of AAV template leading to higher PGI levels and correlation between PGI levels and relative serum hF9 expression levels (Fig. 1c-d). Interestingly, while AAV copy number increased up to 7-fold with higher doses, integration frequency rose by less than 2-fold, suggesting that template DNA is not the primary rate-limiting factor for PGI (Fig. 1e). We then quantified expression in a subset of animals (one from each AAV dose) using a pooled human serum standard and found that the levels of hF9 in animal serum were 20-60% of normal hF9 levels, confirming that the integrated transgene led to functional transcription, splicing, translation, and secretion into the blood at levels that are expected to be curative for patients with F9 deficiency (Fig. 1f).

We then used two sequencing-based methods, short-read linker-mediated PCR (LM-PCR) with the Illumina NGS platform and long-read sequencing with Oxford Nanopore (ONT), to characterize the integration events at the molecular level. We observed uniform template insertion flanked by attL and attR recombination sequences, clear evidence of a deterministic editing event from integrase-mediated PGI (Fig. 1g). Integrase mediated gene insertion is not dependent on generating double stranded DNA breaks (DSBs), and when using circular DNA templates can result in homogenous integration events (Data not shown). Due to the linear nature of the AAV genome, we observed the expected seamless integration on the left side, but on the right side, downstream of the integrated transgene, we detected a series of heterogeneous integration events. This variability is likely caused by inconsistent resolution of the inverted terminal repeats (ITRs) by the host DNA repair machinery. In addition to integrase-mediated integration events, we discovered a significant number of reads showing integration of the entire AAV template in both the forward and reverse orientations (Fig. 1h, Extended data Fig. 2-3). The absence of Bxb1 attL and attR recombination sequences revealed evidence that these genome insertion events were integrase independent. Due to the presence of partial beacon sequences in these reads, we termed these insertions dual flap-mediated PGI (DF-PGI).

**Figure 2.**
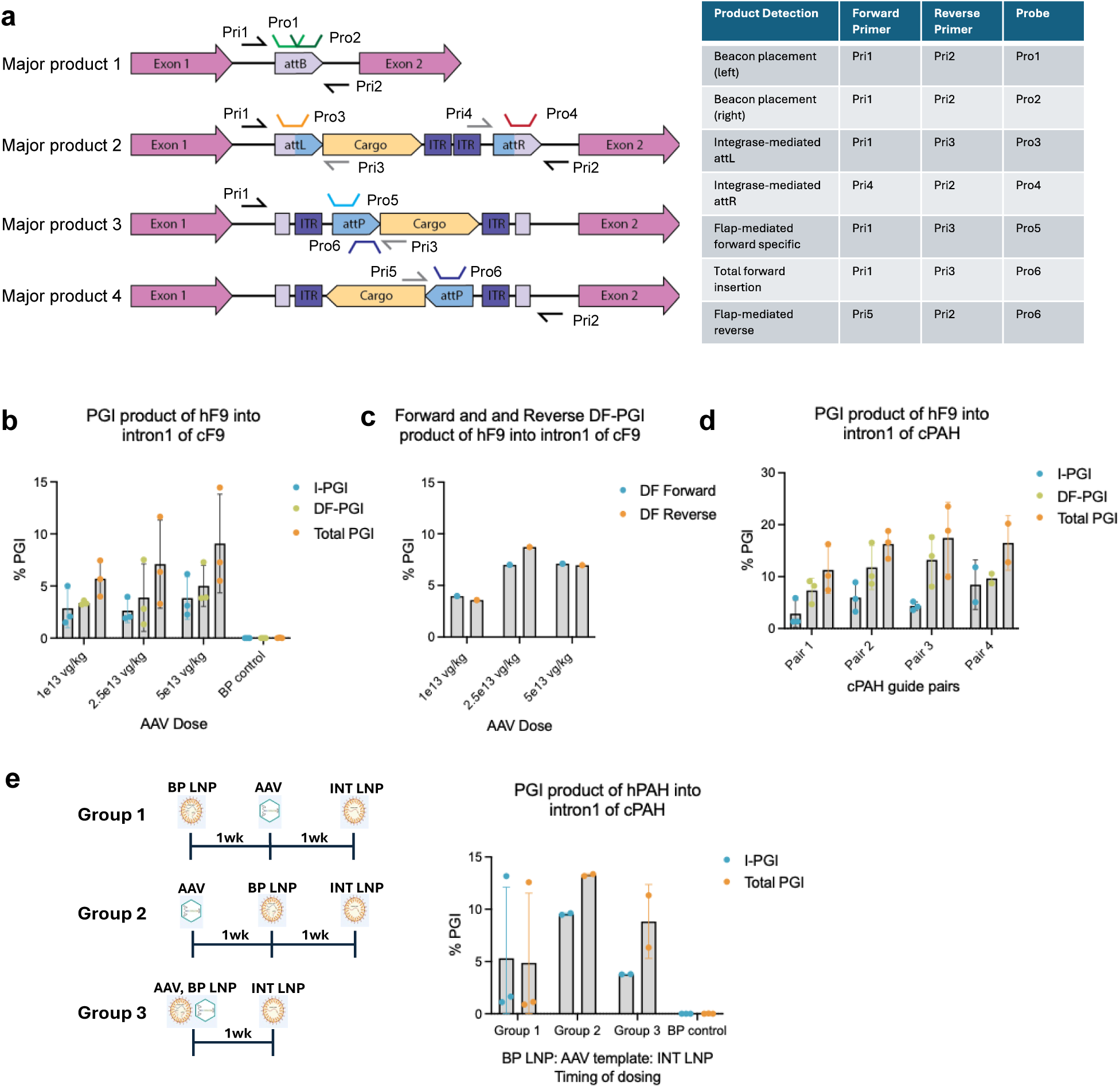
Quantifying PGI outcomes in NHP using digital droplet polymerase chain reaction (ddPCR) (**a**) Generalized graphic depicting all major PGI products resulting from placement of a linear AAV cargo into intron 1 of either F9 or PAH and showing primer and probe placement for detection of each product. Product 1: beacon placement only. Product 2: integrase-mediated AAV integration into placed beacon. Product 3: flap-dependent AAV integration in forward direction with respect to coding sequence. Product 4: flap-dependent integration in reverse direction with respect to coding sequence. All primer and probes labeled as “Pri” and “Pro” respectively with number indicating unique sequences. Table indicates primer and probe combinations for each different assay. (**b**) Measurement of forward PGI frequency at three AAV doses in F9 in NHP by ddPCR. I-PGI was assessed by genomic forward primer, cargo reverse primer, and probe targeting the attL junction. DF-PGI was assessed using the same primers and placing the probe over the left half of the cargo attP which is only present in Bxb1-independent insertion. Total PGI was assessed using the nonspecific cargo probe over the right half of attP. (**c**) Assessment of flap-dependent AAV insertion in the highest efficiency individuals from each AAV dose group in both forward and reverse genomic orientation demonstrating the absence of directional bias in the DF-PGI insertion mechanism. (**d**) Measurement of forward I-PGI, DF-PGI, and total PGI at PAH in NHP by ddPCR showing efficiencies at four different atgRNA guide pairs using the same AAV cargo and mRNAs. (**e**) Dosing schedule diagram for each timing group and DF-PGI versus total PGI frequency in NHP with varying component dose timing. Each group received a different dose schedule with either each component dosed 7 days apart or with co-dosing components staggered with the third component. I-PGI and total PGI frequencies separate (indicating contribution of DF-PGI) only when AAV is dosed before or simultaneously with beacon placement.

**Figure 3.**
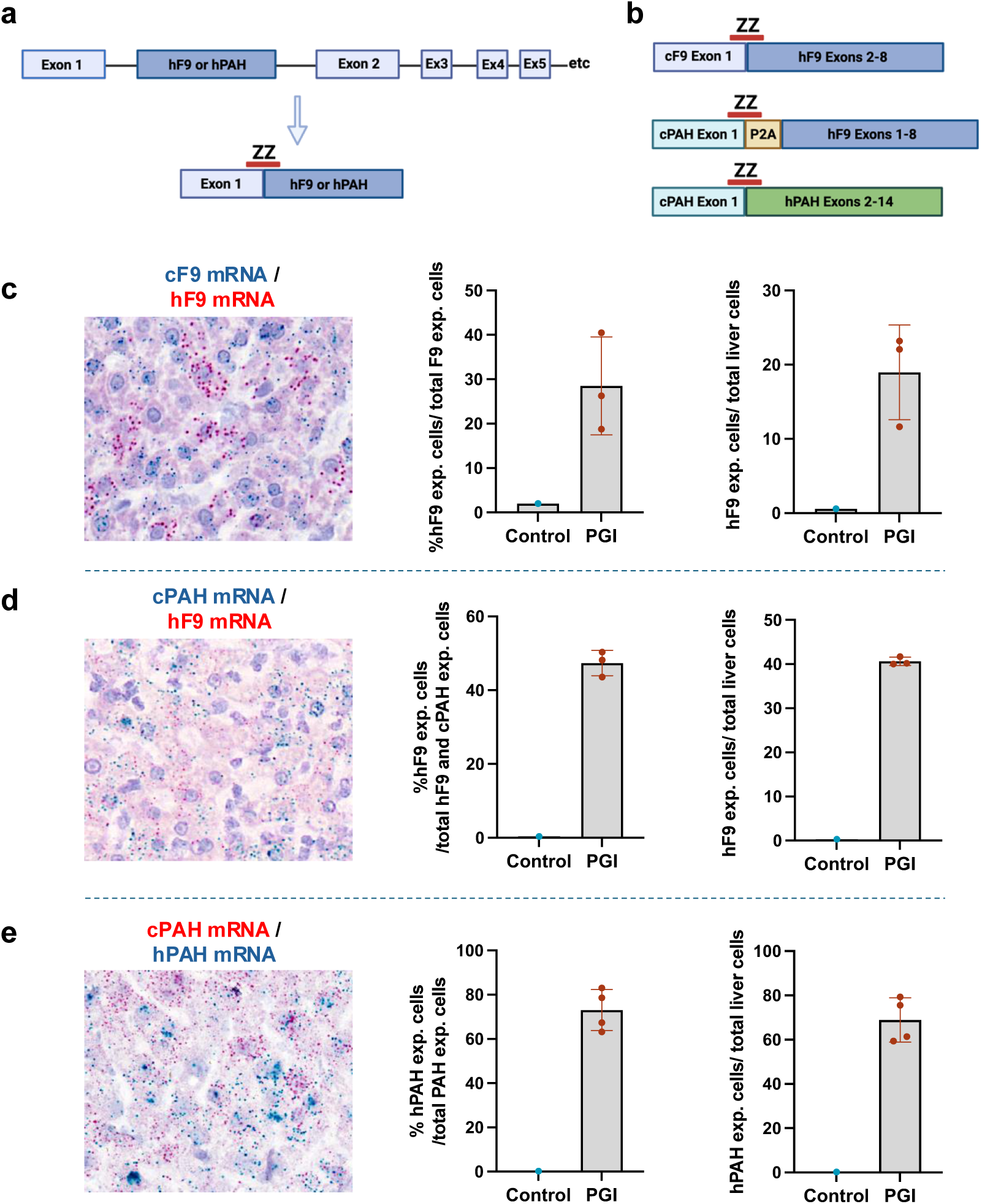
Cellular quantification of PGI efficiency in NHP using BaseScope with three different therapeutic transgene insertions. (**a**) Graphic of Basescope assay showing ZZ probe placement on exon 1 of endogenous gene and exon 2 of transgene allowing specific detection of post-splicing hybrid transcript. (**b**) Schematic representation of probe design for 3 loci and template combinations tested in NHP: hF9 insertion at cF9, hF9 insertion at cPAH, and hPAH insertion at cPAH. (**c**) Representative image of Basescope staining in liver section and cellular quantitation from hF9 insertion at cF9 with two normalization methods. Red signal results from detection of cF9-hF9 PGI transcript and blue signal results from detection of endogenous cF9. HALO quantification of cells expressing fusion mRNA post PGI normalized to cF9 positive only cells or total liver cells. (**d**) Representative image of Basescope staining in liver section and cellular quantitation from hF9 insertion at cPAH with two normalization methods. Red signal results from detection of cPAH-hF9 PGI product and blue signal results from detection of endogenous cPAH mRNA. HALO quantification of cells expressing fusion mRNA post PGI normalized to cPAH positive only cells or total liver cells. (**e**) Representative image of Basescope staining in liver section and cellular quantitation from hPAH insertion at cPAH with two normalization methods. Red signal results from detection of cPAH-hPAH PGI product and blue signal results from detection of endogenous cPAH mRNA. HALO quantification of cells expressing fusion mRNA post PGI normalized to cPAH positive only cells or total liver cells.

AAV ITRs are known to be recognized by DNA repair proteins^9^ and we hypothesized that the host repair mechanism responsible for resolution of flaps generated by RT during beacon writing may also be capable of incorporating AAV genomes. To further investigate this phenomenon, we designed additional ddPCR assays to discriminate between PGI populations (Fig. 2a). Consistent with the sequencing data, we observed that DF-PGI was contributing significantly to total PGI (Fig. 2b). We observed similar DF-PGI frequency in both the forward and reverse orientation as expected based on AAV ITR symmetry (Fig. 2c). This data also strengthens the proposed DF-PGI mechanism as insertion in the reverse orientation is not possible via Bxb1 integrase. This phenomenon was not template or locus specific, as inserting a hF9 or hPAH AAV template into the cyno PAH (cPAH) locus yielded similar results (Fig. 2d-e). Though DF-PGI occurred at all guide pairs tested, the ratio of I-PGI to DF-PGI varied, suggesting that the surrounding sequence impacted DF-PGI efficiency. When we explored component timing, we found that DF-PGI only contributed to total PGI if AAV was pre-dosed or co-dosed with BP-LNP (Fig. 2e). Importantly, high levels of total PGI were achieved in all groups, indicating that both DF and I-PGI can efficiently drive therapeutically relevant levels of targeted gene insertion.

Thus far, we had only measured allelic PGI frequency on bulk liver tissue, which underestimates edit efficiency in hepatocytes due to the presence of other cell types such as Kupffer, stellate, and liver sinusoidal endothelial cells (LSECs). We performed Basescope imaging to determine the number of hepatocytes expressing the hybrid mRNA PGI product (Fig. 3a). Starting with hF9 insertion into the cF9 intron 1, mRNA transcripts for the integrated transgene were detected using a pair of ZZ probes spanning cF9 exon 1 and hF9 exon 2, allowing discrimination between edited and unedited transcripts (Fig. 3b). To preserve NHP life, mouse studies were used to show that the promoterless AAV-hF9 template alone does not drive gene expression (Extended Data Fig. 4a-c). Additionally, probe placement targeting the endogenous exon 1 sequence ensures that only cells with successful PGI are detected (Extended Data Fig. 5a-c).

Basescope quantification (HALO) was performed on the highest efficiency animals from each AAV dose group; 1e13, 2.5e13, and 5e13 vg/kg. We observed hF9 transcript in up to 20% of total cells and 30% of cF9 expressing cells (Fig. 3c). This is aligned with the expected number of edited cells based on bulk total PGI, given that hepatocytes make up 60% of liver cells^10, 11^, the average ploidy of primate hepatocytes is ∼2.5^12^, cF9 is on the X-chromosome, and all animals were male. We observed that most hepatocytes either expressed cF9 (blue) or hF9 (red), with few cells that were double positive due to the transgene preventing endogenous cF9 splicing (Fig. 3c). At the PAH locus, an average of 18% total PGI in bulk liver corresponded to approximately 45% of hybrid mRNA-positive hepatocytes, normalized to the total number of cells expressing hF9 and cPAH (Fig. 2d, Fig. 3d). In contrast to X-linked F9, PAH is on chromosome 12 and is present as two copies per diploid cell. Thus, most hF9-positive cells also expressed cPAH, indicating that the majority of cells had single integration events that allowed the expression of both transcripts.

Phenylketonuria (PKU) is a severe genetic disease that results from mutations in the phenylalanine hydroxylase (PAH) gene^13^. This is one of most common liver-related genetic diseases and while some disease modifying treatments are available for some patients, there is still unmet need for a curative therapy. Given the broad range of mutations that are known to cause PKU (3,370 known associated mutations)^14^, the ideal approach would be integration of a healthy PAH copy into the endogenous locus, which would be curative for most, if not all patients, independent of their individual mutation. To this end, we found that we could efficiently integrate a tagged human PAH cDNA template into the cyno PAH intron1 locus at high efficiency, achieving ∼13% PGI in bulk liver (Fig 2e), corresponding to expression in >60% of liver cells (Fig 3e). These levels of gene insertion are significantly higher than the 10% that is estimated to be curative for PKU^15^, and PGI resulted in protein expression (Extended data Fig. 6). These results were confirmed with human targeting atgRNA in primary human hepatocytes (PHH) using a Flag tagged PAH template. Protein expression is observable after insertion mediated by PGI, with little to no background detected in AAV-only conditions, confirming that our promoterless AAV template alone does not result in gene expression (Extended data Fig. 7).

We further explored the DF-PGI mechanism in primary cynomolgus monkey and human hepatocytes (PCH, PHH). Using transfection timing optimized for total PGI efficiency, we dosed cells with combination treatments to identify the exact variables required for DF-PGI (Fig. 4a-b). We found that as expected I-PGI required nCas9, RT, both atgRNAs, Bxb1, and an AAV template, while DF-PGI was possible under a broad set of conditions. At minimum, DF-PGI required an AAV template and paired nicks, with efficiency significantly increased with at least single flap writing and reached maximum efficiency with paired flaps. Interestingly DF insertion rates with beacon placement only are the same as in full I-PGI conditions, suggesting that AAV opportunistically inserts with partial beacon placement with no impact on Bxb1 activity. Forward and reverse DF-PGI efficiencies were similar for all conditions and the relative integrated protein expression was concordant with total PGI, which was consistent with our observations in NHP (Fig. 4c-d). These data further confirm that I-PGI and forward DF-PGI both result in productive insertion events with the expression of correctly spliced transcripts (Fig. 4e). Thus, DF-PGI is an alternate PGI mechanism that can be combined with I-PGI to result in highly efficient insertion of genes into any genomic location, including their endogenous loci.

**Figure 4.**
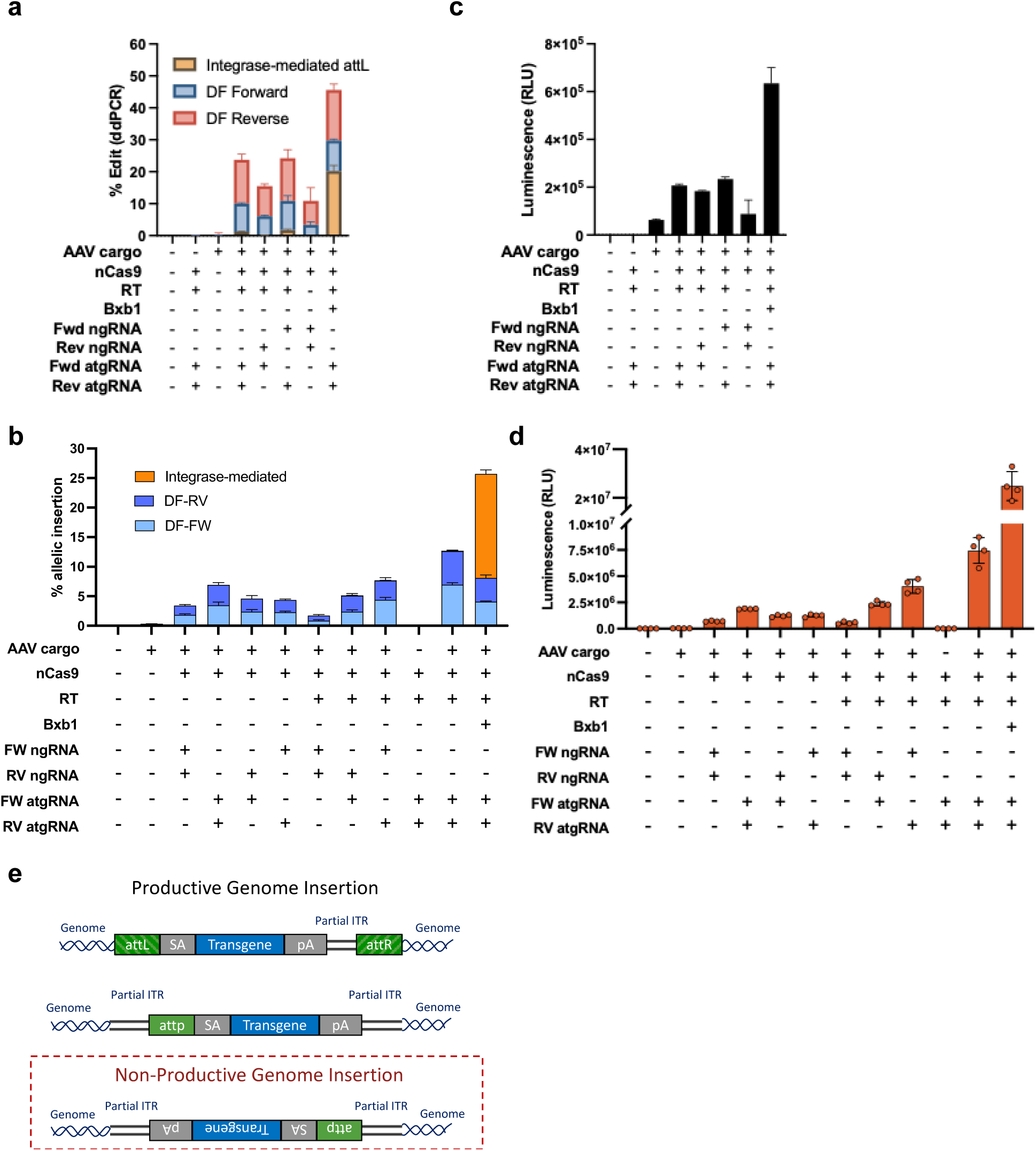
Investigations into DF-PGI mechanism in vitro. In vitro exploration of flap requirements in (**a**) primary cynomolgus monkey hepatocytes (PCH) and (**b**) primary human hepatocytes (PHH). Cells were transduced with AAV 1 day before co-transfection of all other components consisting of combinations of guides and mRNAs as specified per condition. For each species, atgRNA and ngRNA consisted of identical spacers to control for edit location. Efficiencies of each PGI outcome were determined by ddPCR. Relative integrated hPAH protein expression was measured by HiBit luminescence assay in cell culture media after PGI in (**c**) PCH and (**d**) PHH. (**e**) Three unique genome insertion products were generated when AAV template was present during beacon placement with Cas9-RT. I-PGI mediated insertion and the forward orientation of AAV genome captured by non-homologous end joining (NHEJ) DNA repair resulted in productive gene and protein expression. The reverse orientation of AAV genome captured by NHEJ is not a productive genome insertion outcome.

DF-PGI has certain advantages and disadvantages when compared with I-PGI. DF-PGI is dependent on AAV ITRs, unlike I-PGI which can use any dsDNA format (e.g. nanoplasmid, minicircle, hdAd). Unlike I-PGI, DF-PGI is not direction specific, meaning that a unidirectional template would lead to ∼50% unproductive integration events in the reverse orientation. This could be overcome by using a bi-directional template at the cost of further restricting the size of the transgene to ∼1 kb for scAAV and ∼2 kb for ssAAV. DF-PGI also results in the integrated sequence being flanked by heterogenous ITR ‘scars’ because it is dependent on host repair mechanisms, while I-PGI using linear AAV templates results in heterogenous ITRs that are only integrated downstream of the transgene (Extended data Fig. 3). Importantly, I-PGI using a circular DNA template would result in homogenous, directional insertion events that are completely independent of host DNA repair (Data not shown). As a single step process that does not require complete beacon placement or codelivery of integrase, DF-PGI has the potential to be a simpler system than I-PGI and removes the potential risk of integrase-mediated off target integration events^16^.

When compared with Cas9 DSB-mediated AAV template insertion, DF-PGI has the potential to be more efficient as target sites are less likely to be consumed by DSB mediated indels. DF-PGI may also be significantly safer, as it avoids DSBs and would prevent Cas9 mediated off-target integrations or translocations as the probability of two off-target nicks being in close enough proximity to allow for template capture is extremely low.

In summary, we have demonstrated that PGI can be used to efficiently integrate a functional copy of a gene into intron 1 of endogenous genes in NHP liver. To our knowledge, this is the first report of gene replacement under endogenous regulation in an animal. We have also uncovered a novel mechanism, DF-PGI, that can be used in combination with I-PGI to achieve functional gene insertion in >50% of hepatocytes in NHP. Depending on the application, DF-PGI can either be avoided or harnessed by simply changing the dosing schedule. We have demonstrated that both DF and I-PGI are active at multiple genomic target sites, and with multiple template sequences. Integration into intron 1 leads to gene expression from the endogenous promoter, resulting in efficient splicing and translation of the integrated gene. The levels of gene integration resulted in clinically relevant levels of F9 protein expression (average ∼40% normal serum levels) without the need for hyper functional F9 variants (e.g. Pauda^17^) and have exceeded the 10% of hepatocytes predicted to lead to a functional cure for PKU^15^. While we believe there is still significant room for optimization, we have already achieved levels of cellular correction (∼50% of hepatocytes) that are expected to be curative for all recessive monogenic diseases of the liver.

The ability to replace broken genes at the endogenous location at clinically relevant efficiencies using PGI overcomes many limitations of current gene therapy/gene editing systems. PGI allows for endogenous regulation (avoiding over or under-expression), a single product that will treat most if not all patients (independent of mutation), and an edit that will grow with the patient, opening the door to treating pediatric patients before the onset of disease.

## Materials and Methods

### LNP Formulation

LNP formulations containing nucleic acid were formulated by mixing with an ethanolic solution of lipids at a mol ratio of 2.5: 54.2: 32.5: 10.9 (PEG lipid: ionizable lipid: cholesterol: DSPC) with an aqueous solution of nucleic acid in phosphate buffer at offset flow rates (1:4) of lipid to nucleic acid solutions (US11298320B2). LNP were diluted with phosphate buffered saline prior to ethanol removal by tangential flow ultrafiltration, followed by buffer exchange and concentration. The formulations were sterile filtered through a 0.2 μm PES membrane and aliquots were stored frozen at −80°C in Tris-sucrose buffer, pH 8.0.

### mRNA synthesis and purification

In-vitro transcription (IVT) was performed to synthesize mRNA from template plasmid DNA (GenScript) that had been linearized using the BspQI enzyme. Following synthesis, mRNA was purified by tangential flow filtration (TFF), followed by affinity chromatography using POROS^TM^ Oligo (dT)25 Affinity Resin (Thermo Scientific^TM^). Ultrafiltration/diafiltration (UF/DF) was then carried out to concentrate the mRNA the 2 mg/mL and to exchange the mRNA into UltraPure^TM^ DNase/RNase-Free Distilled Water (Invitrogen).

### Cell studies

Cryopreserved primary cynomolgus monkey hepatocytes (MKCP10 CY427, Gibco) were recovered in Hepatocyte Plating Media (A1217601, CM300, Gibco) and plated onto Collagen I coated 96-well tissue culture plates (356698, Corning) at 48k cells per well. Cryopreserved primary human hepatocytes (HMCPMS, Hu8450, Gibco) were recovered in Cryopreserved Hepatocyte Recovery Media (CM7000, Gibco) and plated at 42k cells per well. 8 hours post recovery cells were washed and cultured in maintenance media (A1217601, CM400, A2737501, Gibco). Cells were treated with AAV at 1e6 moi 1 day after plating and transfected with nCas9-RT mRNA, atgRNA 1, atgRNA 2, and Bxb1 mRNA using Lipofectamine MessengerMAX (LMRNA001, Invitrogen) in Opti-MEM (31985062, Gibco) 2 days after plating. Cells were collected for analysis 5 days after transfection.

### Animal studies

Transgenic C57BL/6J mice with a knock-in of the attB site in intron 1 of F9 were generated by Biocytogen (Beijing, China) using CRISPR/Cas9. Mice were transferred to Biomere (Worcester, MA, USA) for breeding. Mice at least 6 weeks old were injected intravenously via tail vein with AAV8 encoding the template DNA at 2e13 vg/kg followed by administration of LNPs containing the integrase mRNA at 3 mg/kg, a week later. All animals underwent euthanasia 7 days after LNP administration and liver lobes were harvested for analysis. All NHP studies used male cynomolgus monkeys (*Macaca fascicularis*) of Cambodian origin were conducted by Altasciences and approved by their Institutional Animal Care and Use Committee.

Animals received LNPs via a 2-hour intravenous infusion at 3mg/kg total RNA. All animals received 2 mg/kg dexamethasone, 0.5 mg/kg famotidine and 5 mg/kg diphenhydramine via intramuscular injection 1 day before treatment, and again 30 minutes before LNP dose administration. Animals received an additional dose of 0.5 mg/kg famotidine, and 5 mg/kg diphenhydramine 1 day post LNP administration. AAV8 was administered by an intravenous bolus injection. All animals underwent euthanasia 14 days after the last dose of test material and liver tissue and serum were collected for analysis.

### gDNA Isolation

Cells were lysed with QuickExtract (QE0905T, Lucigen) and gDNA was purified using SPRI magnetic bead cleanup. Liver tissue was homogenized on Precellys Evolution (cat K002198-PEVO0-A.0 Combo, Bertin technologies, WA, USA). DNA was extracted with quick-DNA/RNA MagBead kit (cat R2131, Zymo research, CA, USA).

### Droplet Digital Polymerase Chain Reaction (ddPCR) Analysis

Custom primers and probes were designed to measure editing in human albumin (hAlb), cyno F9 (cF9), cyno Pah (cPah), and mouse F9 (mF9). Results were normalized to custom reference assays targeting unedited regions of the same genes in the respective species. Probes were dual labelled with 3′-3IABkFQ and either 5′-carboxyfluorescein (FAM) for edit targets or 5′-hexachloro-fluorescein phosphoramidite (HEX) for reference. Assays were validated using gBlocks representing edit outcomes to test for both specificity and linearity. All primers, probes, and gBlocks were synthesized by IDT (Coralville, IA, USA). The reaction mix for all reactions, was composed of 12 µL of 2x ddPCR Supermix for probes (No dUTP) (cat 1863025 Bio-Rad, Hercules, CA, USA), 1.2 µL of each primer and probe mix, 0.12 µL of HindIII (cat FD0505, Thermo Fisher Scientific, MA, USA), 0.12 µL Eco91I (cat FD0394, Thermo Fisher Scientific, MA, USA), 1 µL genomic DNA and water to a final volume of 25 µL. For mRNA quantification from NHP tissue, RNA was extracted from homogenized tissue by Monarch® Total RNA Miniprep Kit (T2010, New England Biolabs). 500 ng DNase I digested RNA was used for cDNA synthesis by Oliog dT primer following the instruction of SuperScript™ IV First-Strand Synthesis kit (18091050, Thermo Fisher Scientific, MA, USA). cDNA was then diluted 100-fold for ddPCR. Droplets were generated on the AutoDG Instrument for automated droplet generation (cat 186410, Bio-Rad, Hercules, CA, USA). PCR amplification was performed with the following cycling parameters: for mice: initial denaturation at 95 °C for 10 min, followed by 40 cycles of denaturation at 94 °C for 30 s and combined annealing/extension step at 58 °C for 1 min, and a final step at 98 °C for 10 min, ending at 4 °C, for NHP: initial denaturation at 95 °C for 10 min, followed by 40 cycles of denaturation at 95 °C for 30 s, annealing at 58 °C for 1 min, extension at 72 °C for 2 min, and a final step at 95°C for 10 min, ending at 4 °C. Data acquisition and analysis were performed on the QX200 Droplet Reader using the QX Manager software (cat 1864003, Bio-Rad, Hercules, CA, USA.)

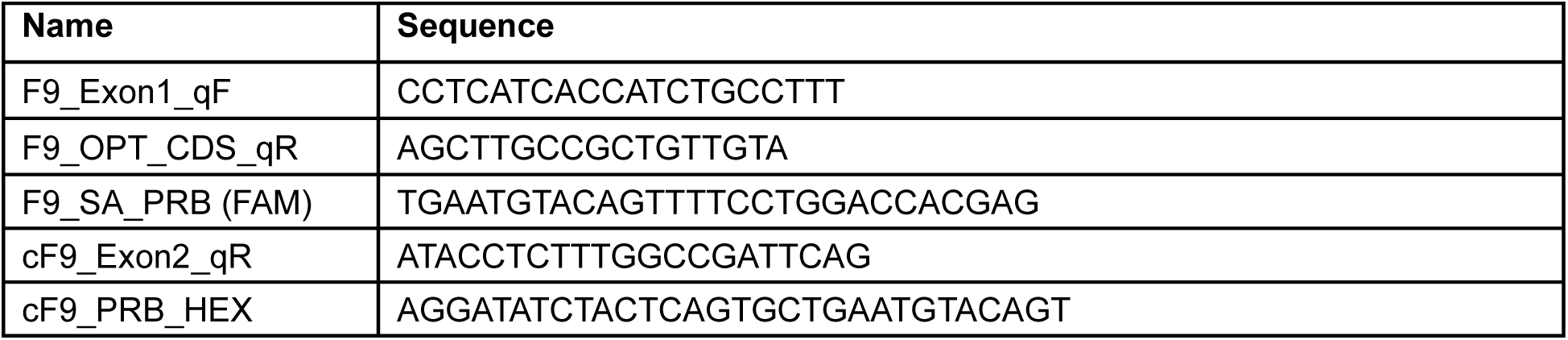

### Oxford Nanopore Sequencing

High molecular weight genomic DNA was isolated from NHP liver tissues using Nanobind Tissue kit from PacBio (Cat. No. 102-302-100) according to manufacturer’s protocol. The concentration of isolated genomic DNA was quantified by broad range Qubit dsDNA quantification assay kit from ThermoFisher Scientific (Cat. No. Q32850). 1µg of isolated genomic DNA was used as a template for PCR amplification to amplify the targeted intronic region of cyno *PAH* gene using the primer set indicated below. PrimeSTAR GXL DNA polymerase from Takara (Cat.No. R050A) was used for carrying out the PCR reaction. PCR amplification was performed with the following cycling parameters: initial denaturation at 98 °C for 3min, followed by 30 cycles of denaturation at 98 °C for 10s, annealing at 60 °C for 15s and extension step at 68 °C for 4.5min, and a final extension step at 68 °C for 3min. Purification of PCR amplicons are carried out using Qiagen QIAquick PCR & Gel Cleanup Kit (Cat. No. 28506) according to manufacturer’s protocol. The purified PCR amplicons were used to generate Oxford Nanopore Technology (ONT) libraries for performing long read sequencing using ONT Native Barcoding Kit 23 V14 (Cat. No. SQK-NBD114.24). The ONT libraries were generated according to manufacturer’s protocol. In brief, end prep reaction was carried out for each samples using 200 fmol of purified PCR amplicons followed by native barcode ligation. For the last step of library preparation, ONT adapter was ligated to the pooled barcoded samples. 10 fmol of the final pooled ONT library was loaded onto MinION Flow Cell R10.4.1 from ONT (Cat. No. FLO-MIN114) and the sequencing run was carried out for 2 days. The long read sequencing data obtained from ONT was analyzed using the in-house bioinformatics pipeline.

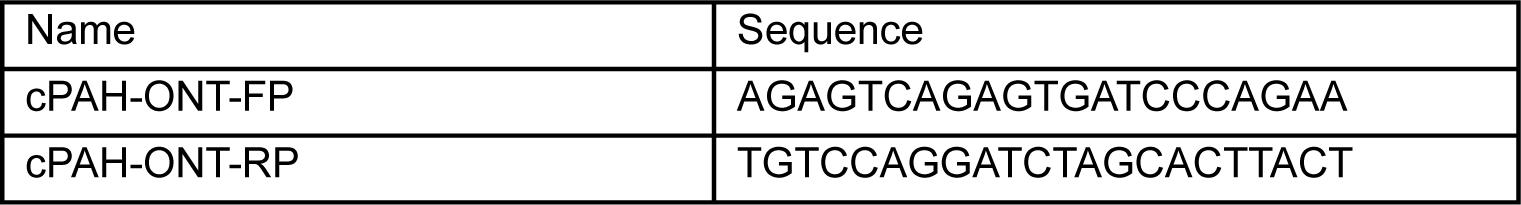

### Data analysis for Nanopore data

Nanopore duplex reads containing both 5’ and 3’ adapters and longer than 2500bp were selected for further analysis. The filtered reads were mapped to prebuilt I-PGI and DF-PGI reference sequences using the Minimap2 aligner^18^. The resulting alignments were visualized in IGV^19^ to confirm the correct structures of I-PGI and DF-PGI.

### Linker-Mediated Polymerase Chain Reaction (LM-PCR)

Genomic DNA samples were prepared for NGS library preparation via LM-PCR by tagmentation of 300ng input DNA with Tn5 transposase recombinant enzyme (GenScript) to achieve desired fragment size of 400-600 bp and to incorporate 5’ overhangs containing patterned unique molecular identifiers (UMIs). Tagmented fragments were enriched and indexed for NGS over two rounds of LM-PCR using NEB Q5 polymerase system (cat M0494S) with a nested gene-specific primer approach. Indexed libraries were sequenced on a MiSeq™ System (Illumina) at a minimum 1 million paired-end reads per sample.

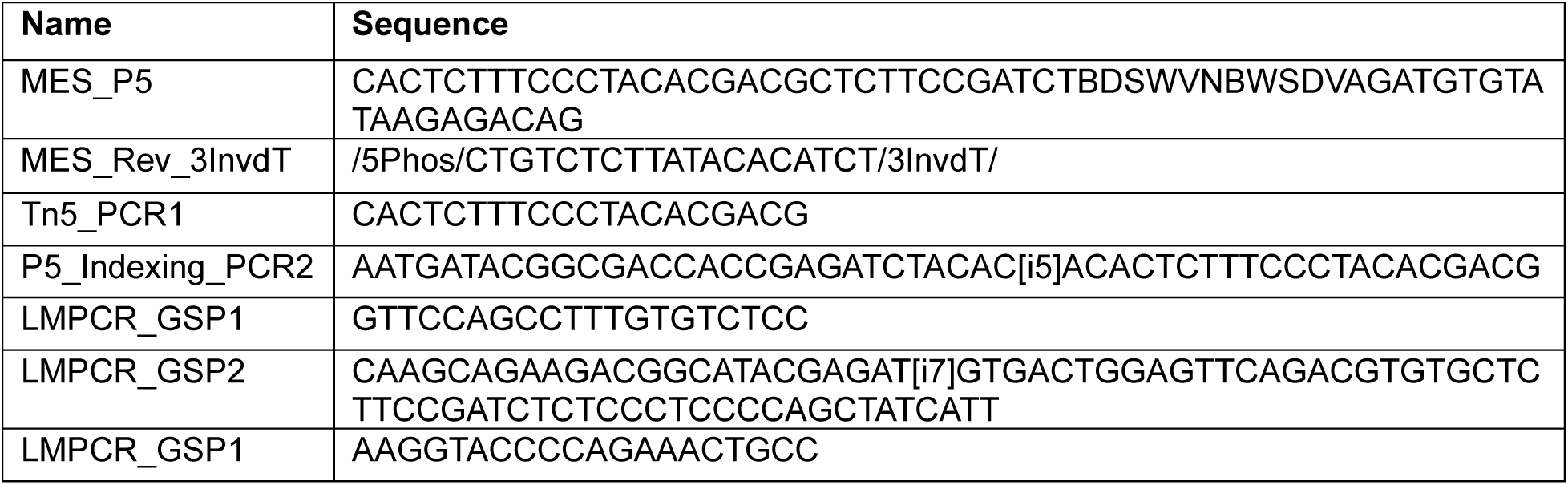
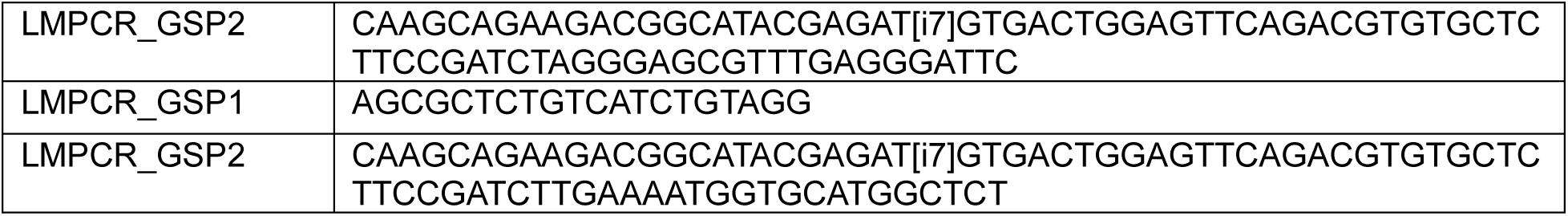

### Sequencing Data analysis for LM-PCR data

Code related to LM-PCR NGS analysis is publicly available on GitHub: https://github.com/jessie-wangjie/tbREVEAL. Briefly, 11-nucleotide (nt) unique molecular identifiers (UMIs) were extracted and removed from Read 1 (R1) raw sequencing reads. The Tn5 adapter in R1 and genome-specific adapter in Read 2 (R2) were trimmed using cutadapt^20^, allowing a maximum of two mismatches. Reads containing both adapters were mapped to the reference genome and cargo reference using aligner BWA^21^. Picard^22^ was employed to mark duplicated reads based on the extracted UMI and mapped position. To predict potential translocations, the split-read detection tool fgsv^23^ was used. Translocations with one breakpoint at the expected genomic insertion site were retained for downstream quantification. Translocations with the second breakpoint at the attP site in the cargo reference were classified as I-PGI, while the second breakpoint at the cargo reference outside of attP were designated as DF-PGI. For each translocation, the corresponding read sequences were extracted and subjected to multiple sequence alignment using MUSCLE^24^. A random subset of these aligned reads was selected for inclusion in the figures.

### HiBiT quantification

HiBiT from integrated PAH cargo was detected using the Nano-Glo HiBiT lytic detection system (N3050, Promega, Madison, WI, USA) following manufacturer protocol. For cell studies, 50 uL of master mix containing 100:2:1 ratio of Nano-Glo HibiT Lytic Buffer, Substrate, and LgBiT Protein was added per cell well and incubated for 10 minutes while shaking and read after 10 minutes additional incubation using GloMax Explorer (cat GM3500, Promega, Madison, WI, USA) with 0.3s integration. For in vivo studies, 125 uL of master mix was used for 25 µL of serum with all other steps identical. For detecting hPAH-HiBiT, liver tissue was homogenized in Nano-Glo HibiT Lytic Buffer on Precellys Evolution (cat K002198-PEVO0-A.0 Combo, Bertin technologies, WA, USA). Lysate was diluted 1:320 in Nano-Glo HibiT Lytic Buffer and 25 µL of the latest was mixed with 125 uL of master mix. All other steps are identical as previously described.

### ELISAs

Human F9 in mouse serum was quantified by ELISA using hF9 ELISA kit (cat LS-F22871, Lifespan Biosciences, Lynnwood, WA, USA) following the manufacturer’s protocol. Mouse serum was diluted 10x in Reference Standard & Sample Diluent, provided by the kit. Expression of hF9 was normalized to human serum.

To detect human F9 in NHP serum, Reacti-Bind 96-well microplates were coated with 100 µL/well (cat PI15041, VWR, PA, USA) of capture antibody mouse mAB to human Factor IX antibody (cat AHIX-5041Haematologic technologies, VT, USA) at a concentration of 3 μg/ml in PBS. Plate was incubated overnight at 4 °C. Plate was washed 3 times with 300 µL/well with ultrapure water (cat 10977-015-023, Thermo Fisher Scientific, MA, USA). Blocking was performed with 100 µL of chonblock (cat 50-152-6971, Thermo Fisher Scientific, MA, USA) for 1 hour at RT. Wells were washed with 300 µL with 1XPBS-0.05% Tween 20. Human serum (cat 092931149, CELLect human pooled serum, MP biomedicals, OH, USA) and monkey serum (cat IGMNCY SER50ML, Innovative research, Inc, MI, USA) were mixed 1:1 as the highest standard. The standard was diluted 1:2 in monkey serum for a total of 7 standards. All standards and sample serum were further diluted 1:100 in PBS. Plate was loaded with 50 µL of standards and sample serum and incubated for 1 hour at RT. Wells were washed three times with 300 µL with 1XPBS-0.05% Tween 20. 50 µL per well of Sheep anti-human Factor 9 polyclonal antibody at 100 ng/mL (cat ab128048, Abcam, MA, USA) were added to each well and plate was incubated for 1 hour at RT. Wells were washed three times with 300 µL with 1XPBS-0.05% Tween 20. 50 µL per well of Donkey anti-Sheep IgG pAbs with HRP at 100 ng/mL (cat ab97125, Abcam, MA, USA) were added to each well and plate was incubated for 1 hour at RT. Wells were washed three times with 300 µL with 1XPBS-0.05% Tween 20. 75 µL of working solution of TMB Substrate Reagent set (cat 555214, BD Biosciences, MA, USA) were added to each well and plate was incubated in the dark for 15 mins. 75 µL of stop solution (cat SS04, Invitrogen, MA, USA) were added and optical density was read at 450 nm.

### BaseScope

RNA in situ hybridization assays were performed using the Basescope Duplex assay (cat 323810, Advanced Cell Diagnostics) following the manufacturer’s protocol. Briefly, the tissues were treated with peroxidase hydrogen blocker before boiling at 98°C–100°C in target retrieval for 10 min. Paired double-Z oligonucleotide probes designed to specifically detect the junction of exon 1 of cynomolgus monkey and exon 2 of human transgene mRNA (Supplemental Figure 5) (BaseScope probe; Advanced Cell Diagnostics) were hybridized for 2 h at 40°C, followed by a series of signal amplification and washing steps. All the incubation steps at 40°C were performed in a HybEZ Hybridization System. Hybridization signals were detected by chromogenic reaction using FAST Red and Green dyes. Slides were counterstained with Gill’s hematoxylin. Bright-field images were acquired using a 3D-Histech scanner with Slide Viewer Software using a 40× objective. Each sample was quality controlled for RNA integrity with a probe specific to the housekeeping genes PPIB and UBC (BaseScope Positive control probes, cat 703191 and 727021-C2, Advanced Cell Diagnostics). Negative control background staining was evaluated using a probe specific to the bacterial dapB gene (BaseScope negative control probe, 700141, Advanced Cell Diagnostics). Quantitative image analysis was conducted using HALO^®^ ISH software (v4.1.3) (Indica Labs, Albuquerque, NM). This analysis software provides cell by cell quantitative results based on number of dots per cell which is categorized into 5 bins (Bin 0 = 0 dots/cell, Bin 1 = 1 dots/cell, Bin 2 = 2-3 dots/cell, Bin 3 = 4-10 dots/cell, Bin 4 = >10 dots/cell). Each punctate dot represents a single mRNA molecule. For each sample, endogenous and transgene mRNA expression levels were analyzed. Analysis settings were optimized for the detection of cells and for each mRNA target.

### PGI of Primary Human Hepatocytes

Frozen primary human hepatocytes were rapidly thawed at 37C, then poured into 50 mL Hepatocyte Thaw Medium (Invitrogen, #CM7000). The cells were centrifuged at 100g for 10 minutes. The supernatant was removed and then the cells were resuspended in 10 mL Hepatocyte Plating Medium via gentle rocking (Invitrogen, #CM3000). Cells were counted, then seeded at 42k cells/well into a 96-well collagen I coated platen in a final volume of 100 µL (Corning, 152038). Cells were allowed to settle evenly for 5-10 minutes before incubating at 37C, 5% CO2 for 6 hours. Plating media was then removed and replaced with 100 µL of maintenance media composed of Williams E Medium + Maintenance Medium Supplement (Invitrogen #A1217601, #CM4000). 24 hours after plating, cells were washed once with maintenance medium. Cells were then treated with AAV-LK03 containing a human PAH coding sequence that was codon optimized and includes a C-terminal 3xFlag tag (MOI of 1e6). 24 hours after AAV cells were washed then treated with maintenance medium containing atgRNAs (4 pmol), nCas9-RT mRNA (2 µg), bxb1 mRNA (2 µg), and 100 µM dTTP/dGTP. After 48 hours, media was replaced with fresh maintenance medium. 72 hours later, cells were harvested for ddPCR and immunocytochemistry staining.

### Immunocytochemistry Staining

Medium was removed and cells were washed once with PBS, then fixed for 10 minutes at room temperature with a 4% Paraformaldehyde-PBS solution (Thermo Fisher #28906; Gibco #10010023). Cells were washed 3x with PBS, then permeabilized with a PBS + 0.1% (v/v) solution of Triton X-100 (ThermoFisher #HFH10) for 10 minutes at room temperature. Cells were washed 3x with PBS, then blocked with a solution of PBST (PBS + 0.05% Tween® 20 and 5% (v/v) normal goat serum) for 1 hour at room temperature (Fisher Scientific #BP337-500; ThermoFisher #PCN5000). Blocking solution was removed, then cells were treated with 100 uL of anti-Flag monoclonal antibody (Cell Signaling, #33542S) diluted 1:200 in PBST. Plate was incubated at 4C overnight. The next day, cells were washed 4x with PBST, then Incubated at room temperature protected from light with a 1:1,000 dilution of Goat anti-Mouse IgG (H+L) Cross-Adsorbed Secondary Antibody, Alexa Fluor™ 594 (2μg/ml final concentration diluted in blocking buffer; ThermoFisher #A-11005). Cells were then washed 4x with PBS before staining with CellBrite® Green Cytoplasmic Membrane Dye as per manufacturer instructions (Biotium, #30021). Cells were washed 3x with PBS, then nuclear staining was performed via DAPI dye for 10 minutes (ThermoFisher R37606). After a final 3x wash with PBS, images were acquired via fluorescence microscope (EVOS m5000). Quantitative image analysis was conducted using a combination of CellProfiler^25^ v4.2.6 to process images and automatically count nuclei, followed by manual quantification of red fluorescing cells by four operators. The percentage of PAH+ cells was calculated as the number red fluorescing PHH cells out of total number of nuclei within each image. The total number of nuclei was adjusted to account for multinucleated PHH cells assuming 20% multinucleation.

### Data analysis

Data were processed using Microsoft Excel and GraphPad Prism 10.0 software packages.

---

**Extended Data Figure 1.**
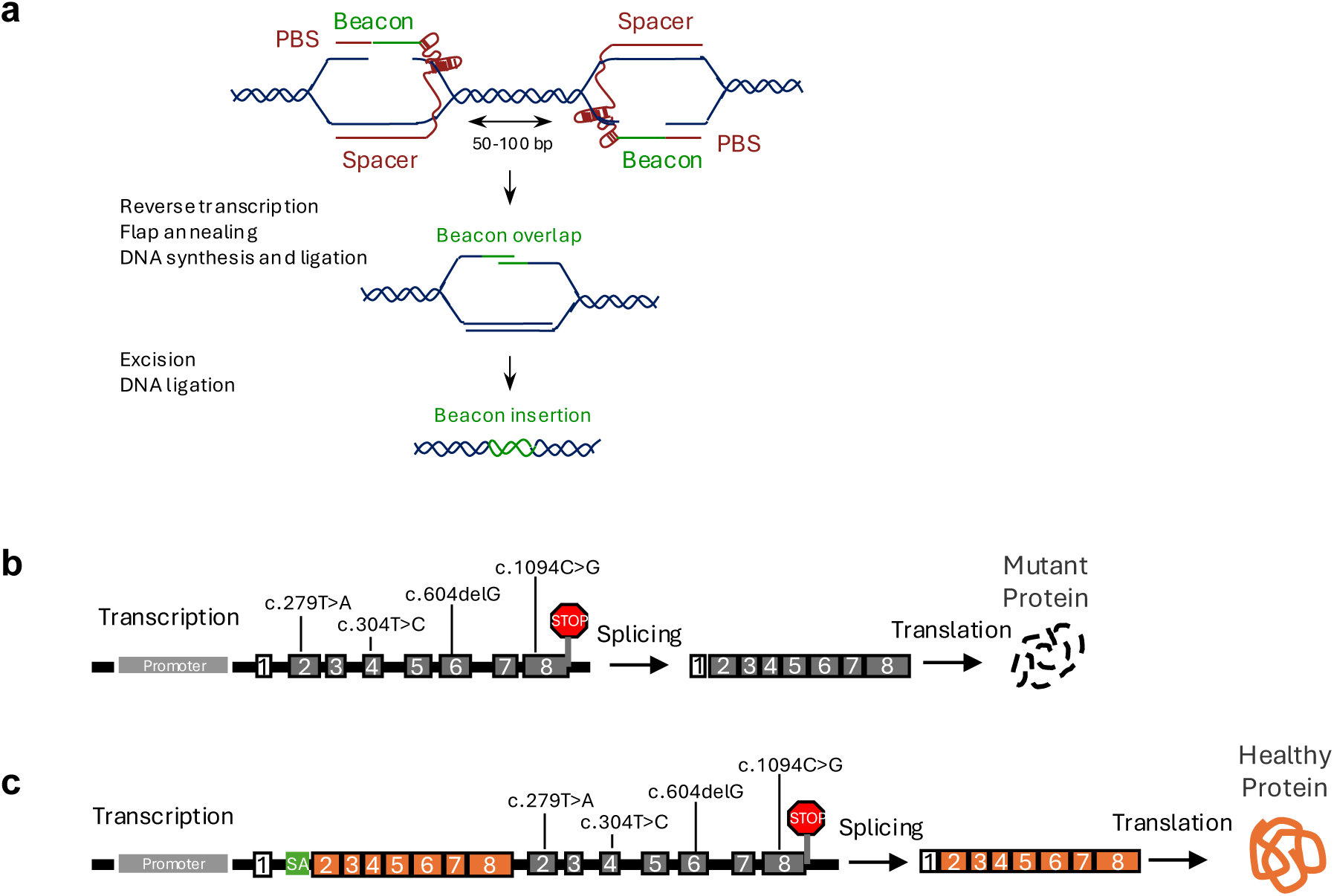
Integrase-mediated programmable insertion (I-PGI) for therapeutic gene replacement is a two-step process that allows all mutations downstream of exon 1 to be corrected with the same drug. (**a**) Two attachment guide RNAs (atgRNAs) target opposing PAMs separated by 50-100 bp. Each gRNA encodes 29 bp of the 38 bp beacon with 20 bp of complementary overlap. After nCas9 makes a nick in the genome, reverse transcriptase copies the partial beacon sequences, which allows complementary base pairing between the 2 DNA flaps with beacon sequence. Endogenous DNA repair promotes excision of the endogenous sequence, gap filling DNA synthesis of the remaining beacon sequence, and ligation to permanently install the beacon into the host genome. (**b**) Diagram depicting diverse pathogenic genetic mutations detected across the gene body from a population of hemophilia B patients which result in mutant protein. (**c**) Using I-PGI, a template containing splice acceptor, healthy copy of downstream coding DNA sequence, and stop codon is inserted into intron 1 that leverages the endogenous promoter to produce healthy gene transcripts for protein translation capable of rescuing gene function for all mutations.

**Extended Data Figure 2.**
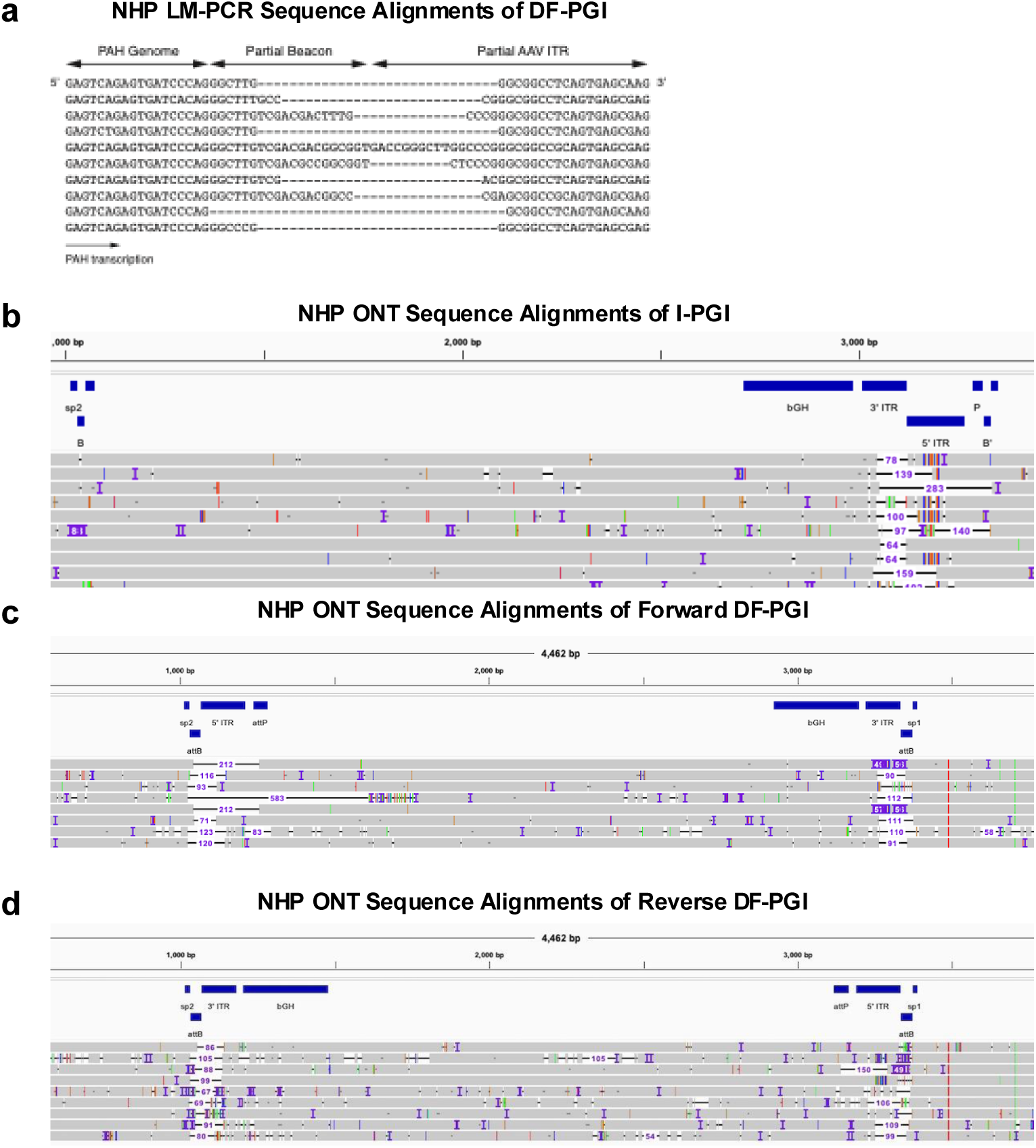
Characterization of PGI editing outcomes with LM-PCR NGS and Long-Read Sequencing with Oxford Nanopore (ONT) (**a**) LM-PCR NGS alignments of DF-PGI reveal partial ITR and partial beacon sequences at the targeted nicking location in cF9 intron 1. (**b**) Representative long-read sequencing alignments of I-PGI reveal small deletions at the location of DNA repair after LSI recombination of linear cargo creates a DSB that requires repair across a new 5’ and 3’ ITR junction. (**c**) Representative long-read sequencing alignments detected in the forward insertion direction for DF-PGI reveal small deletions at the genome ITR junctions. (**d**) Representative long-read sequencing alignments detected in the reverse insertion direction for DF-PGI reveal small deletions at the genome ITR junctions.

**Extended Data Figure 3.**
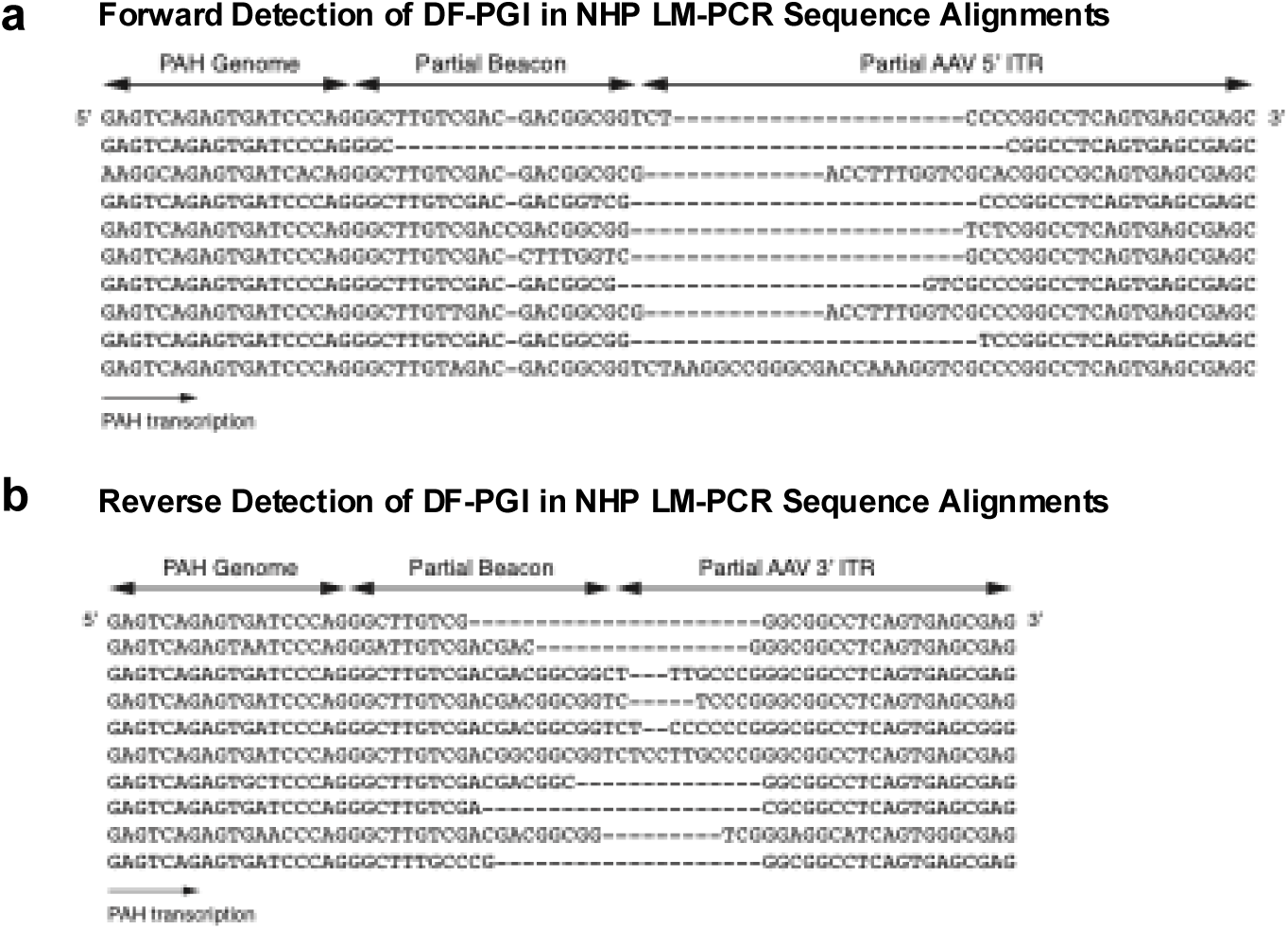
Detection of bidirectional insertion via DF-PGI with LM-PCR. (**a**) LM-PCR NGS alignments of DF-PGI detect genome junctions associated with the 5’ ITR. (**b**) LM-PCR NGS alignments of DF-PGI also detect genome junctions associated the 3’ ITR, which is an indication of bidirectional insertion.

**Extended Data Figure 4.**
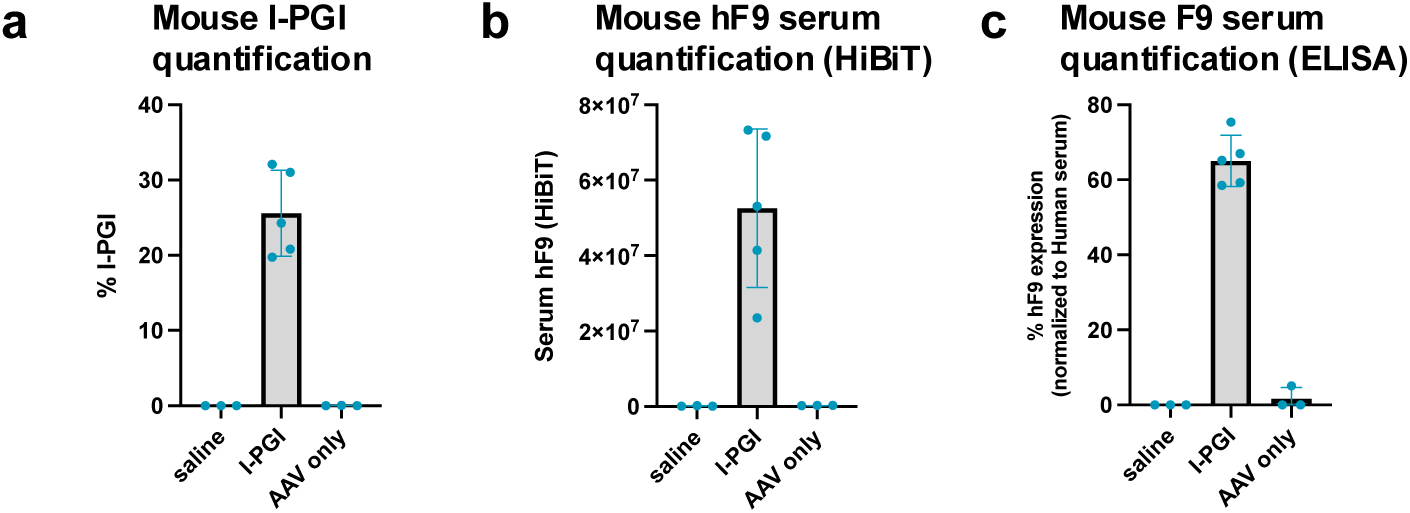
Quantification of I-PGI in male F9-attB mice. (**a**) %I-PGI of hF9 template integration into attB at mouse F9 (mF9) by ddPCR. (**b**) Relative quantification of mouse serum hF9-HiBiT using luminescence reporter assay. (**c**) Quantification of mouse serum hF9 against pooled human serum standard using ELISA. Data shows successful I-PGI in group injected with AAV and integrase (I-PGI group) and no genomic integration or protein expression detected in serum of mice injected with AAV only.

**Extended Data Figure 5.**
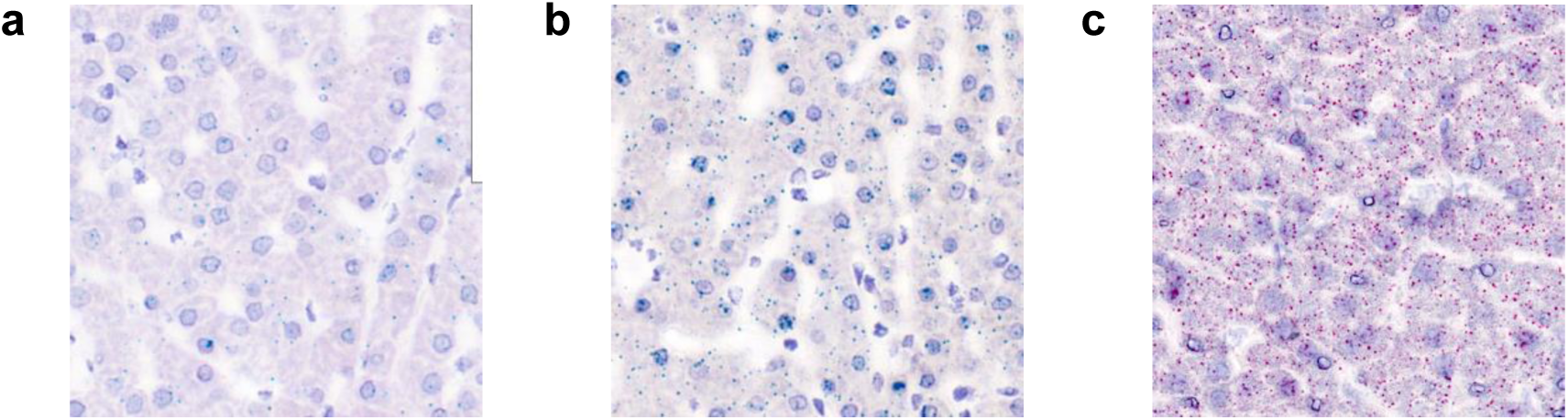
BaseScope images of untreated control NHP livers. (**a**) ACD Bio pre-qualified liver: cF9 transcript labeled blue, hybrid cF9-hF9 transcript labeled red. (**b**) ACD Bio pre-qualified liver: cPAH transcript labeled blue, hybrid cPAH-hF9 transcript labeled red. (**c**) Liver from untreated NHP: Endogenous cPAH transcripts labeled red, hybrid cPAH-hPAH transcripts labeled blue. No background staining was observed for hybrid transcripts.

**Extended Data Figure 6.**
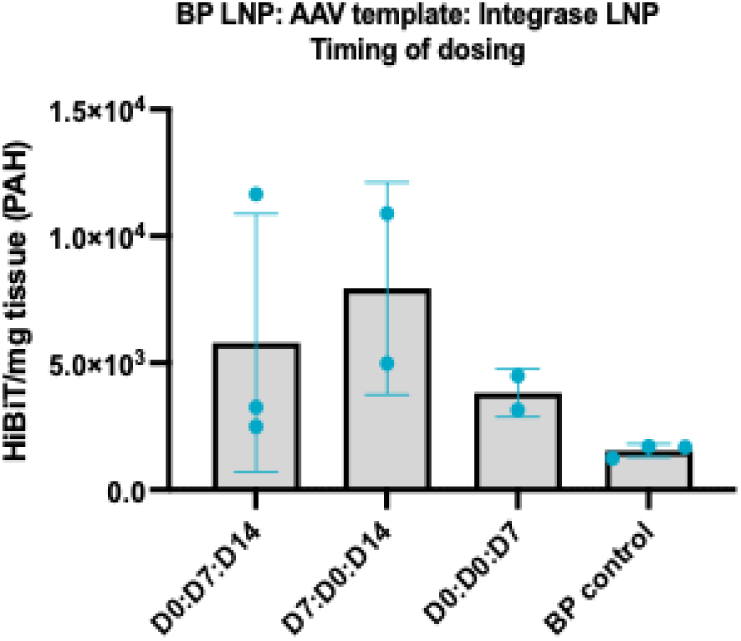
Quantification of hPAH protein expression post PGI in NHP. Relative hPAH levels in tissue lysate from insertion of the HiBit tagged gene determined by luminescence assay.

**Extended Data Figure 7.**
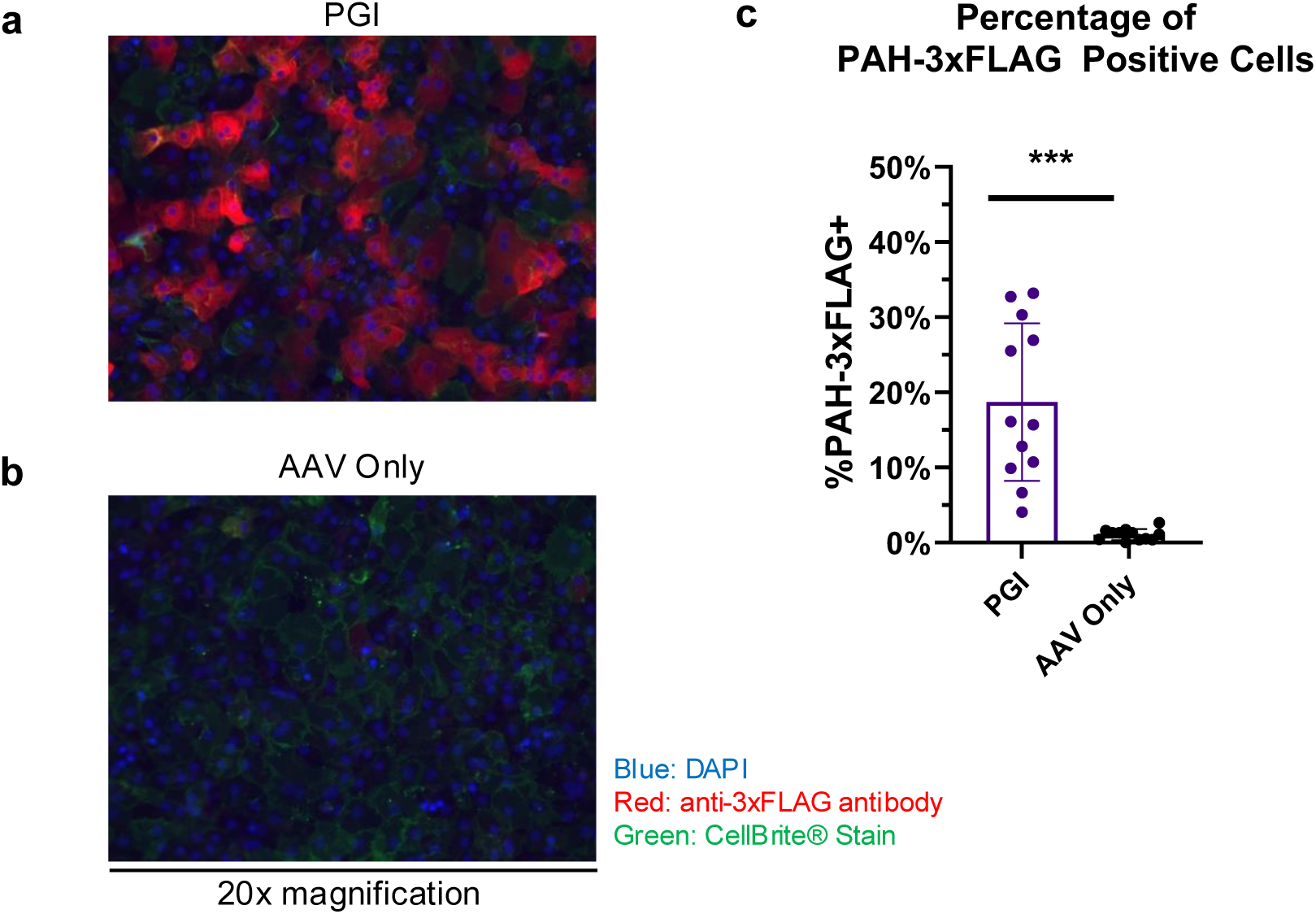
PAH-Flag3x Immunocytochemistry staining of Primary Human Hepatocytes that have undergone PGI at the PAH locus. (**a**) Representative images at 20x magnification of the positive staining detection of PGI transgene in PHH in contrast to control samples (**b**) treated with AAV only. Red channel: anti-3xFLAG antibody, Blue channel: DAPI nuclei stain, and Green Channel: CellBrite® Cell Membrane Stain. (**c**) Raw images were processed through CellProfiler software and the percentage of PHH cells with positive staining (PAH-Flag+) was estimated by dividing by the total number of nuclei in each image and assuming 20% of PHH were multinucleated.

## Notes

### Competing Interest Statement

The authors have declared no competing interest.

